# Evolutionary insights into provirus-encoded CRISPR-Cas systems in halophilic archaea

**DOI:** 10.1101/2025.06.17.660064

**Authors:** Doron Naki, Uri Gophna

## Abstract

Prokaryotic microorganisms coexist with mobile genetic elements (MGEs), which can be both genetic threats and evolutionary catalysts. In *Haloferax lucentense*, a halophilic archaeon, we have recently identified an unusual genomic arrangement: a complete type I-B CRISPR-Cas system encoded on a mega-plasmid coexists with a partial counterpart within an integrated provirus in the main chromosome. The provirus-encoded system lacks the adaptation genes (*cas1, cas2*, and *cas4*), suggesting its potential reliance on the plasmid-encoded CRISPR-Cas module for the acquisition of new spacers. This arrangement suggests a potential instance of “adaptive outsourcing,” where a provirus might leverage a co-resident MGE for a key function. Through comparative genomics, we show that similar proviral CRISPR-Cas systems are found in distantly related haloarchaea (e.g., *Natrinema* and *Halobacterium*), indicating probable virus-mediated horizontal transfer and suggesting they may function as mobile defense modules. Phylogenetic analysis highlights distinct evolutionary origins of the two systems: the plasmid system clusters with other *Haloferax* CRISPR-Cas systems, while the proviral system clusters with those from other genera, consistent with horizontal acquisition. Interestingly, spacer analysis reveals that the proviral systems predominantly target viral sequences, while the plasmid system appears to target both plasmids and viral sequences, a distribution mirroring broader trends observed in other plasmid- and chromosome-encoded CRISPR systems. This observed targeting preference suggests a potential for complementarity that could support a model of cooperative immunity, where each system may protect its genetic “owner” from competition and, indirectly, the host.

## Introduction

Archaea and bacteria coexist with a diverse community of mobile genetic elements (MGEs), ranging from plasmids and transposons to viruses of varied morphologies (Luk et al. 2014). Interactions with these MGEs often result in a dynamic molecular arms race, wherein both hosts and MGEs continually evolve strategies of attack and defense (Koonin et al. 2020a). While MGEs are often regarded as parasitic, accumulating evidence suggests they can also confer valuable traits, fueling genetic innovation and adaptation in their hosts (Koonin et al. 2017; Rocha & Bikard 2022; Al-Shayeb et al. 2022). Specifically, horizontal gene transfer driven by MGEs has enabled prokaryotes to acquire novel functions—from antibiotic resistance in bacteria to virus resistance in archaea—and hence contribute broadly to microbial evolution (Ochman et al. 2000; Baltrus et al. 2013).

Within this evolutionary context, CRISPR-Cas (Clustered Regularly Interspaced Short Palindromic Repeats and CRISPR-associated proteins) systems have emerged as a central adaptive immune strategy in prokaryotes (Barrangou et al. 2007; McGinn & Marraffini 2019; Hille & Charpentier 2016; Barrangou 2013a). These loci store short “spacer” sequences acquired from invading foreign DNA, thus allowing the host cell to recognize and neutralize related genetic elements upon subsequent encounters (Sorek et al. 2013). In archaea, CRISPR-Cas systems display remarkable diversity, with various subtypes (I, II, III, IV, V, VI) (Koonin et al. 2017b) and an array of organizational architectures (Makarova et al. 2015; Makarova et al. 2011). Certain archaeal species even harbor multiple, functionally distinct CRISPR-Cas loci, hinting at overlapping or modular forms of adaptive immunity (Turgeman-Grott et al. 2019). Although CRISPR-Cas modules are typically encoded chromosomally, they have also been identified on plasmids and integrated viral genomes (proviruses), underscoring the complex gene exchanges that characterize archaea–MGE interactions (Pinilla-Redondo et al. 2022). Furthermore, some CRISPR-Cas systems, particularly Type IV and some Type VI, frequently lack adaptation modules and are thought to rely on trans-acting Cas1/Cas2 proteins from other systems (Pinilla-Redondo et al. 2022; Silas et al. 2017), demonstrating that modularity and “outsourcing” of some CRISPR functions is not uncommon.

*Haloferax lucentense*, a halophilic archaeon, exemplifies this genetic and ecological complexity. Like other species in the *Haloferax* genus, it inhabits hypersaline environments where viruses and plasmids abound (Luk et al. 2014). We have recently revealed an unusual genomic configuration in *H. lucentense*: a type I-B CRISPR-Cas system fully encoded on a self-replicating plasmid, coexisting alongside a partial type I-B CRISPR-Cas locus within a 36 kb provirus. Notably, the provirus-encoded system lacks the canonical adaptation genes (*cas1, cas2, cas4*), thereby rendering it incapable of independently acquiring new spacers. Such “incomplete” or “defective” CRISPR-Cas modules have been described in other archaea and bacteria (Al-Shayeb et al. 2022), yet the confluence of proviral and plasmid-encoded systems in a single host, particularly with evidence of functional interaction, is less well understood.

Theoretically, harboring dual CRISPR-Cas systems could strengthen host immunity by broadening target specificity or by creating multiple interference layers. Alternatively, certain models—such as the Black Queen Hypothesis—suggest that organisms can offload metabolically expensive or potentially risky functions onto co-residents of the same niche (Morris et al. 2012; Van Valen et al. 1973). In the context of *H. lucentense*, this perspective implies that the provirus may “outsource” its adaptation needs to the plasmid’s full CRISPR-Cas repertoire while maintaining its own interference capabilities. This arrangement, that we termed “adaptive outsourcing,” could enable the host to benefit from an additional layer of defense provided by the provirus, while the provirus, by maintaining interference capabilities, could effectively repel additional MGEs, including competing viruses or invasive plasmids. This scenario could confer a tangible selective advantage to both the host and the integrated virus. (Al-Shayeb et al. 2022). The provirus, by shedding its adaptation genes, may avoid the risk of self-targeting that could arise from uncontrolled Cas1/Cas2 activity. Alternatively, the smaller CRISPR-Cas cluster may be beneficial in the context of a compact viral genome that often is limited by the need to fit into a small capsid.

In this study, we dissect the genomic, phylogenetic, and functional features underlying these dual CRISPR-Cas systems in haloarchaea. By integrating long- and short-read sequencing data, we present the complete organization of the proviral and plasmid-encoded loci in *H. lucentense*. We then analyze such systems in diverse haloarchaea with particular emphasis on spacer content, interference genes, and adaptation modules. Furthermore, we illustrate how these systems exhibit complementary targeting preferences—suggesting a division of labor in recognizing plasmids versus viruses, consistent with broader genomic surveys—and explore how partial CRISPR arrays remain effective via “adaptive outsourcing.”

## Results

### Diverse haloarchaeal proviruses encode type I-B CRISPR-Cas systems

*H. lucentense* harbors a unique configuration of CRISPR-Cas systems: a complete Type I-B system on a plasmid and an incomplete Type I-B system within a provirus integrated into the main chromosome. Both systems are classified as type I-B, but the provirus-encoded system lacks the adaptation module, presumably relying on the plasmid-encoded system for acquiring new spacers.

This unusual arrangement raises questions about the origins, functions, and evolutionary implications of having such dual systems. To obtain a complete genome sequence of *H. lucentense*, we employed a hybrid sequencing approach utilizing both Illumina and Nanopore sequencing technologies (Jain et al 2016). Longer reads generated from Nanopore sequencing were particularly valuable for resolving repetitive regions and complex genomic architectures. The resulting data facilitated a hybrid assembly of the *H. lucentense* genome and its plasmids through the Trycycler assembly pipeline.(Wick et al. 2021b) This approach not only allowed us to differentiate between the main chromosome and the plasmids but also enabled the identification of the circular replicating form of the provirus, which is only rarely shed into the growth medium. Further details on the assembly strategy are provided in the Materials and Methods section.

### Comparative Genomics of Proviral CRISPR-Cas Systems

To further investigate the evolutionary history and functional potential of the provirus-encoded CRISPR-Cas system in *H. lucentense* (family Haloferacaceae), we conducted a comparative analysis with related proviruses. This included comparing it to proviruses found in other *Haloferax* strains and more distantly related Haloarchaea, as illustrated in Figure 1. Our synteny analysis revealed a high degree of conservation in gene content and order between these *Haloferax* proviruses, particularly in regions encoding essential viral functions such as DNA replication, packaging, and structural components, functions which are typical for such viruses. (Turgeman-Grott et al. 2024). However, a striking difference was observed in the presence of the CRISPR-Cas system. While the *H. lucentense* provirus harbors a type I-B CRISPR-Cas system, this system is absent in the corresponding regions of the related *Haloferax* proviruses. This finding suggests that the CRISPR-Cas system was acquired horizontally after the *H. lucentense* provirus diverged from other viruses in this family. The acquisition of this CRISPR-Cas system may have provided a selective advantage, by enabling CRISPR-Cas targeting of competing viruses or other MGEs.

**Figure 1.**
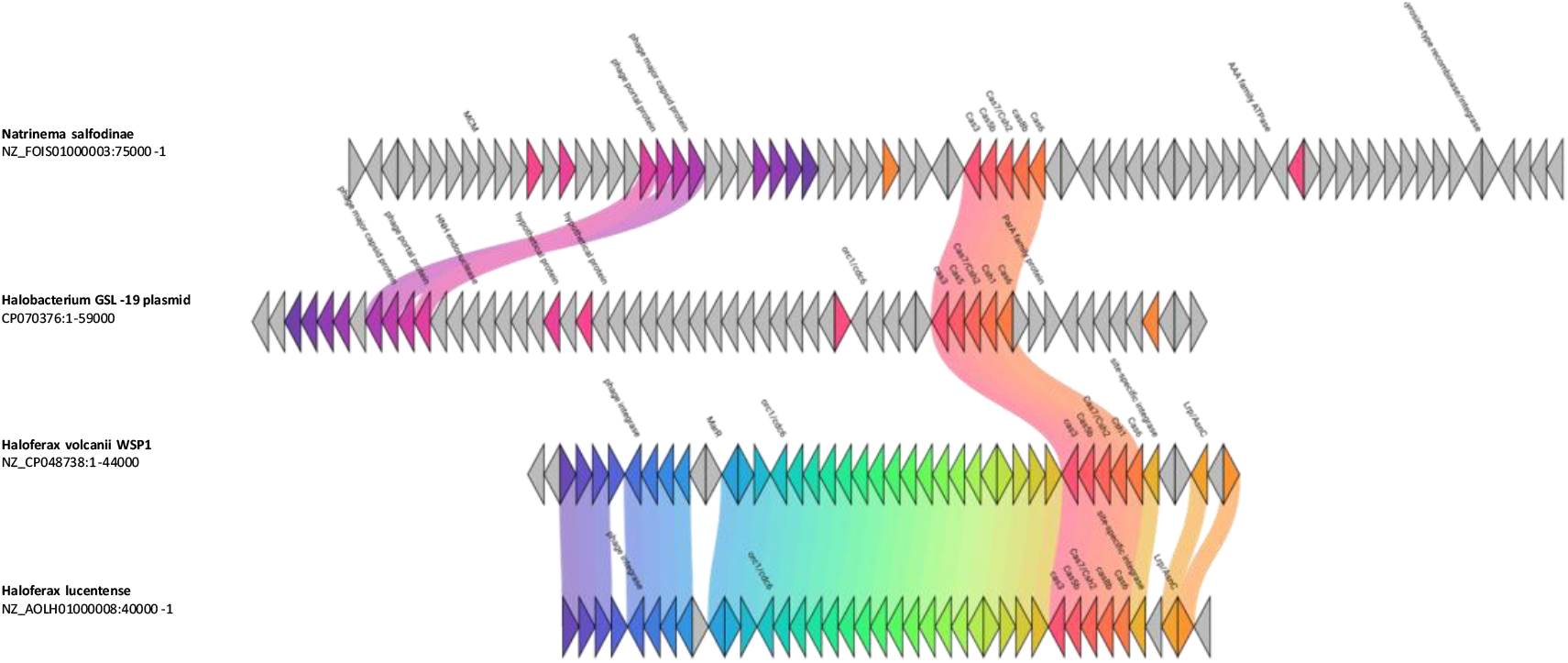
Comparative gene maps of haloarchaeal viruses carrying partial CRISPR-Cas systems. Synteny diagrams show the gene organization of viral genomes that encode an adaptation-deficient type I-B CRISPR-Cas module (as in Haloferax lucentense). Colored arrows denote ORFs (color-coded by predicted function or orthology), and shaded bands connect homologous regions with >75% amino acid identity (identified by clinker (Gilchrist et al. 2021)). The H. lucentense proviral CRISPR-Cas locus is highly similar to those observed in the proviruses from Natrinema and Halobacterium, despite otherwise having completely dissimilar genomes. This conservation of the CRISPR-Cas region across different viruses suggests horizontal transfer of a defense module among haloarchaeal viruses.

Subsequent comparative genomics analysis revealed that CRISPR-Cas systems highly similar to the one within the *H. Lucentense* provirus (70-90 % amino acid identity) are found in viruses infecting distantly related Haloarchaea, specifically *Natrinema salifodinae* (family Natrialbaceae) and *Halobacterium* GSL-19 (family Halobacteriaceae). Notably, unlike *H. lucentense*, in both *N. salifodinae* and *H*. GSL-19, these adaptation-deficient systems represent the sole CRISPR-Cas locus identified within their respective genomes. These systems are situated within distinct provirus remnants that show little similarity to the *H. lucentense* provirus. Furthermore, the context varies: while in *Natrinema salifodinae* the CRISPR-encoding provirus region is in the main chromosome, in *H*. GSL-19, the system resides in a provirus locus within a 54.9 Kb plasmid that harbors four CRISPR arrays. This widespread distribution across families and genomic contexts strongly suggests that these provirus-associated CRISPR-Cas systems are capable of horizontal transfer across different archaeal virus lineages. Furthermore, as shown in Figure 1, the proviruses that harbor these nearly identical CRISPR-Cas systems (e.g., in *H. lucentense, Natrinema salifodinae*, and the one within the *Halobacterium* GSL-19 plasmid) can be otherwise largely dissimilar in their genomic content, highlighting the modularity of the CRISPR-Cas region itself.

The presence of such similar, yet genomically isolated, CRISPR-Cas systems in distant relatives implies they confer a significant selective advantage, potentially enabling the viruses to compete with other viruses and additional MGEs as well as potentially contribute to host defenses, aligning with the “guns for hire” concept (Koonin et al. 2020a). This observation resonates with findings of other virus-encoded CRISPR-Cas systems in bacteria, including streamlined versions potentially reliant on host factors (Al-Shayeb et al. 2022). In *H. lucentense*, the proviral element encodes the interference machinery but “outsources” spacer acquisition, a dependency potentially mirrored in the *Natrinema* and *Halobacterium* systems given their lack of adaptation modules.

In contrast to these divergent contexts, *Haloferax* sp. WSP1 presents a case of remarkable conservation. This strain, isolated in India (Verma et al. 2020), harbors both a provirus and a plasmid-encoded CRISPR-Cas system that are highly similar to those found in *H. lucentense* (isolated in Spain). Comparative analysis revealed extensive spacer identity between the two strains. Specifically, twenty-one spacers from the plasmid arrays of *H. lucentense* are shared with the plasmid arrays of WSP1, all of which currently have no identifiable targets. Additionally, two spacers from the proviral arrays of *H. lucentense* are shared with the WSP1 proviral system, also with unknown targets. Most notably, one spacer from the *H. lucentense* plasmid array, which targets the internal membrane protein of HFPV1 (*Haloferax volcanii* pleomorphic virus 1) (Alarcón-Schumacher et al. 2022), is also present in the WSP1 plasmid array. This pattern of shared spacers, spanning both systems and geographic distances, suggests either recent horizontal exchange or strong stabilizing selection in response to conserved MGE threats across disparate environments. Whether these shared spacers confer functional immunity to HFPV1 in both strains warrants experimental investigation.

### Spacer Content Analysis Reveals Expanded Targeting, Functional Specialization, and Widespread Immune Memory Against Dominant Haloarchaeal Viruses

A comprehensive analysis of 50 unique spacers derived from the CRISPR arrays of haloarchaeal genomes encoding type I-B systems of viral origin (25 arrays from 9 genomes) revealed a complex and specialized targeting landscape. By querying spacers against the NCBI RefSeq viral, IMG/VR and RefSeq Plasmid databases(Sayers et al. 2023; Paez-Espino et al. 2017), 84 target matches (out of 805 spacers) were identified across 37 distinct genomes, expanding the known network of mobile genetic element (MGE) interactions in these haloarchaea.

Targeting Patterns were affected by system origin: spacers from plasmid-associated CRISPR systems predominantly targeted plasmids and phage-derived elements, consistent with a defensive role against horizontally transmitted plasmids. In contrast, provirus-associated spacers were enriched for matches against viral genomes and prophages, suggesting specialization in interviral competition. Solitary chromosomal arrays contributed a smaller but ecologically meaningful number of antiviral spacers. These system-specific trends are visualized in Figure 2.

**Figure 2.**
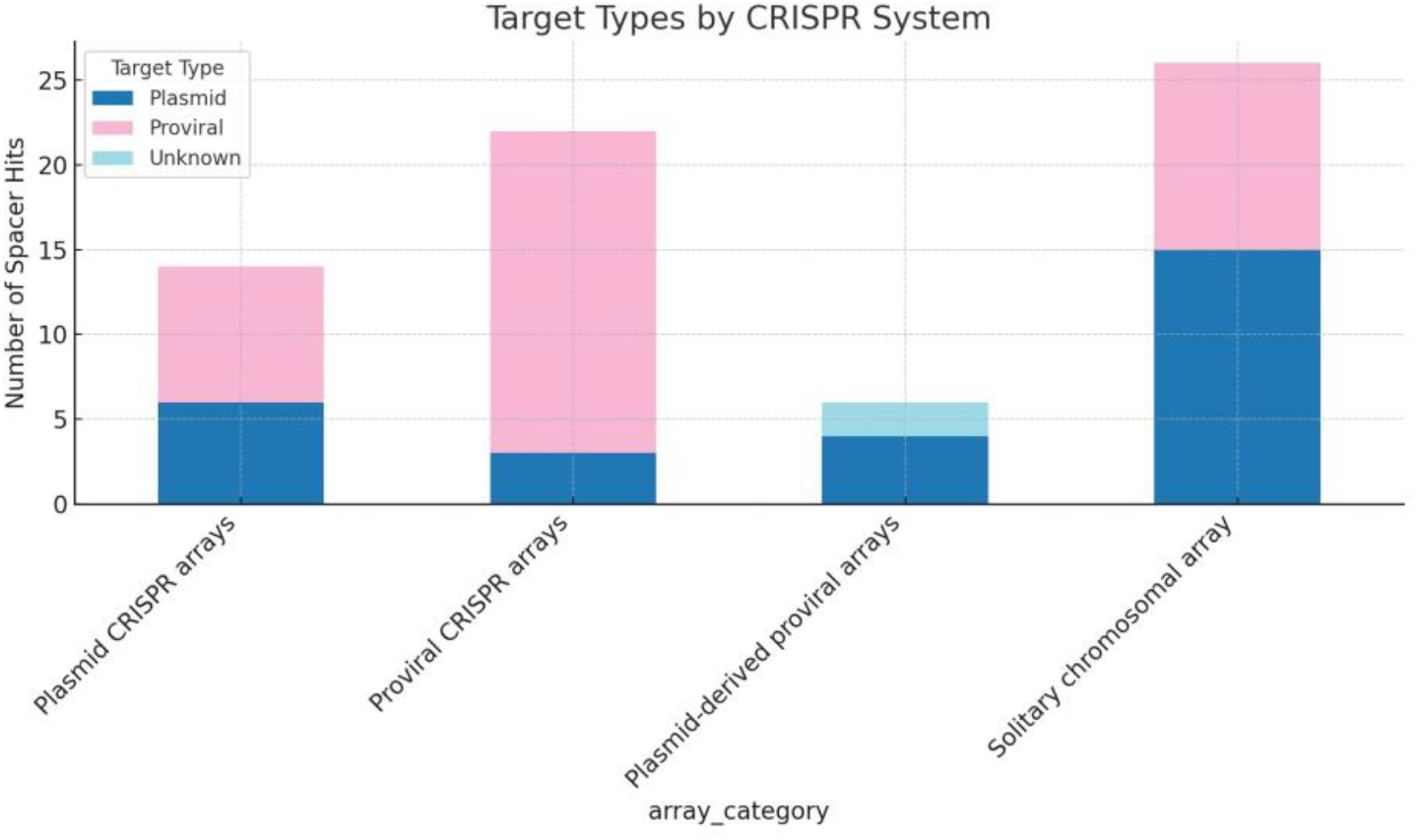
Spacer target preferences of plasmid vs proviral CRISPR-Cas systems in Haloferax. Bar plots showing the number of CRISPR spacers (grouped by their source locus) matching three target categories: plasmid, viral (including proviruses), or unknown sequences. Each bar corresponds to a CRISPR array from H. lucentense viral/plasmid system or related systems. The targeting profiles differ by system: plasmid-encoded CRISPR systems target plasmids (blue bars) and proviruses(pink bars), while provirus-encoded systems overwhelmingly target viruses (pink bars), with chromosomal (solitary) arrays contributing to both type of targets. These patterns highlight a division of labor – plasmid CRISPR-Cas immunity is geared towards plasmid and viral threats, whereas proviral CRISPR-Cas primarily counters viral competitors – together providing a broad defense.

Curiously, a notable finding was that spacers targeting two previously studied viruses that infect *Haloferax*, HFPV1 (*Haloferax volcanii* pleomorphic virus 1) and LSV-48N (Lemon shaped virus of *Haloferax* strain 48N) were frequently observed (Alarcón-Schumacher et al. 2022; Turgeman-Grott et al. 2024). HFPV1 was targeted by seven independent spacers originating from *H. lucentense, Haloferax* sp. BAB-2207, and two additional *H. volcanii* strains (SS0101 and wsp1). Targets included internal membrane proteins (the majority), as well as the spike protein, and several hypothetical proteins, suggesting broad recognition of essential viral components. LSV-48N was targeted by two independent spacers, one from *H. lucentense* and one from *H. volcanii* strain wsp1, both directed against a hypothetical protein. The presence of multiple, independent targeting spacers against both HFPV1 and LSV-48N across different CRISPR arrays and strains suggests that these viruses, or at least closely-related ones, represent common threats encountered by these haloarchaeal hosts, thereby maintaining selective pressure for CRISPR-Cas mediated immunity against them.

Beyond viral targets, multiple plasmids were independently recognized by several spacers. *Haloferax volcanii* plasmid pHV3 (438kb), *Haloferax volcanii* plasmid pSVX82-1 (577kb), *Haloferax volcanii* plasmid pBNX82-1 (540kb) and *Natrinema pallidum* plasmid pNMX15-1 (56kb), indicating acquisition of immunity against prevalent mobile genetic elements. None of these plasmids encode CRISPR-Cas.

Beyond individual targeting events, the data also point towards potential interplay between the two systems. Notably, we observed instances where plasmid- and provirus-derived spacers from the same organism targeted nearby regions on the same viral genome (*Haloferax* Atlit-12N and Atlit-6N proviruses), a finding that strongly suggests the potential for cooperative immune action against MGEs. Another interesting observation was a subset of spacers from *H. volcanii* wsp1 arrays that targeted helix-turn-helix transcriptional regulators encoded on plasmids. This specialization likely reflects selective pressure to disrupt plasmid replication and stability mechanisms. Generally, most spacers targeted viral structural proteins (spike proteins, tail-length tape measure proteins), transcriptional regulators (helix-turn-helix domains, PadR family repressors), DNA-related proteins (Zn-ribbon binding proteins, RecT recombinases, and restriction-modification methylases), suggesting inter-MGE conflicts. These results demonstrate a modular, cooperative CRISPR immune network shaped by shared ecological pressures across *Haloferax* strains.

### Phylogenetic Analysis Reveals Distinct Evolutionary Origins and Horizontal Transfer of Proviral- and Plasmid-Encoded CRISPR-Cas Systems

Comprehensive phylogenetic analysis of Cas proteins from both viral and non-viral systems provides compelling evidence for their distinct evolutionary origins. The plasmid-encoded system clusters closely with CRISPR-Cas systems found in various *Haloferax* species (mostly encoded on megaplasmids), suggesting it represents the canonical type I-B system widely distributed throughout this genus. In contrast, the provirus-encoded system appears sporadically across different genera of Haloarchaea, demonstrating its propensity for horizontal transfer. Phylogenetic trees constructed using Cas6 sequences clearly separate the two systems into distinct clades; the proviral Cas6 and its cohorts cluster with sequences from *Natrinema* and *Halobacterium* viruses rather than with the host’s plasmid-encoded Cas6, consistent with horizontal transfer across genera of viruses and hosts (Figure 3). This phylogenetic separation, combined with the differing GC content (63.4% vs 66.8%), strongly supports an independent evolutionary origin for the proviral systems.

**Figure 3.**
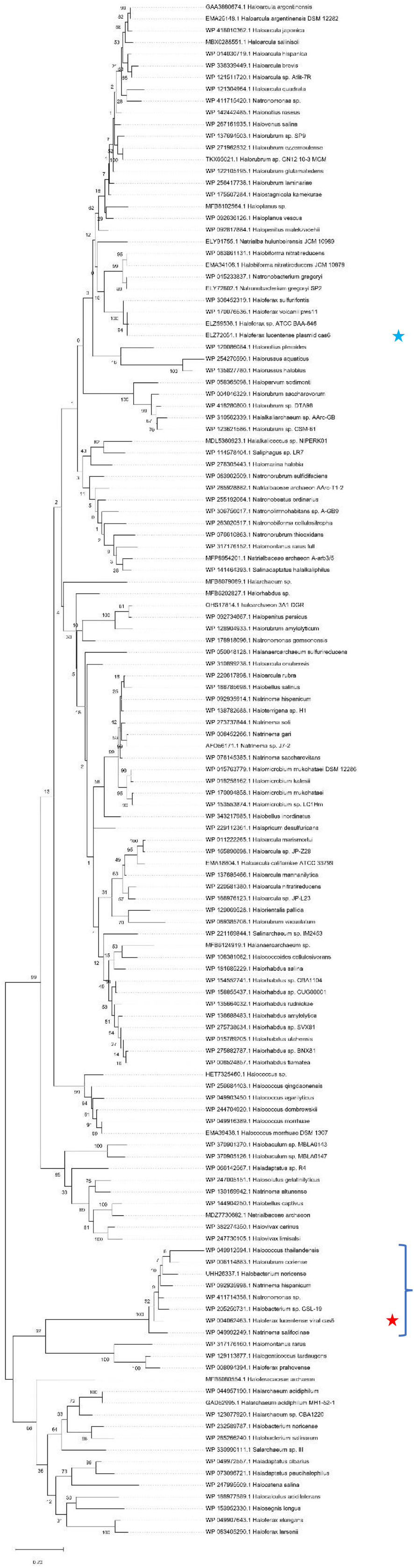
Phylogenetic separation of virus-encoded vs plasmid-encoded Cas6 proteins. A maximum-likelihood phylogenetic tree of 142 Cas6 endoribonuclease sequences from diverse haloarchaea showing two distinct clades. The H. lucentense provirus-encoded Cas6 (marked with a red asterisk) clusters with Cas6 proteins from other archaeal viruses, whereas the plasmid-encoded H. lucentense Cas6 (blue asterisk) groups with Cas6 from other Haloferax species. Bootstrap support values (100 replicates) are indicated at key nodes. The clear split between viral and plasmid Cas6 lineages indicates independent evolutionary origins: the proviral Cas6 was likely acquired horizontally (from a viral gene pool), while the plasmid Cas6 reflects a more vertical evolution with transfers within the Haloferax genus. (Tree constructed using a JTT+Gamma model implemented in Mega12; scale bar denotes 0.2 amino acid substitutions per site.)

Intriguingly, while the *cas* genes of related proviruses showed high sequence conservation (>75% nucleotide identity), the remaining viral genes exhibited substantial divergence. This unusually high similarity suggests a relatively recent horizontal acquisition of the *cas* gene cluster in these viruses.

Phylogenetic trees for other Cas proteins (Cas7, Cas5, etc.) show a similar grouping (supplementary_data_1).

### Complete CRISPR-Cas Systems Phylogenetically Proximal to the Proviral Clade Exhibit Signatures of Mobility

The Cas6 phylogenetic tree (Figure 3) revealed a distinct clade of provirus-encoded Cas6 proteins. Interestingly, a small sister clade to these proviral systems included Cas6 proteins from complete CRISPR-Cas systems found in *Halomontanus rarus* KZCA124, *Haloferax prahovense* DSM 18310, and *Halegenticoccus tardaugens* SYSU A00711. Given their phylogenetic position and completeness (i.e., possessing adaptation modules), we investigated their genomic contexts for signatures of mobility that might link them to the evolutionary trajectory of proviral systems.

In *Halomontanus rarus* KZCA124, the complete Type I-B system (including *cas1b, cas2, cas3’’, cas4, cas5b, cas6, cas7b, cas8b*) is encoded on a plasmid (NZ_CP135524). Notably, this plasmid-borne system is flanked by genes indicative of mobilization: a tyrosine-type recombinase/integrase gene is located ∼1.2 kb upstream of *cas6*, and an IS5 family transposase gene is present ∼6.3 kb further upstream. The synteny of this region is detailed in Supplementary Figure S1. Similarly, the complete Type I-B CRISPR-Cas system in *Haloferax prahovense* DSM 18310, located on contig NZ_AOLG01000031, is also associated with MGE-related genes. An ISH3 family transposase (pseudo) is found ∼200 bp downstream of the *cas3’’* gene, and a site-specific integrase is located ∼1.5 kb downstream. Upstream of the CRISPR array, ParA and Orc1/Cdc6 family replication proteins are also present (Supplementary Figure S1).

The complete Type I-B system in *Halegenticoccus tardaugens* SYSU A00711 (contig NZ_SDIC01000002) presents a slightly different context. While lacking immediate flanking transposases or integrases within a 10kb upstream window, an RNA-guided endonuclease of the TnpB family is located ∼2 kb downstream of the *cas3’’* gene. TnpB proteins are themselves frequently associated with mobile elements (Supplementary Figure S1).

The association of these complete CRISPR-Cas systems, which are phylogenetically close to the proviral Cas6 clade, with plasmids and/or MGE-associated genes (integrases, transposases, TnpB) supports the hypothesis that such mobile, yet complete, systems could be the evolutionary precursorto the partial, adaptation-deficient CRISPR-Cas systems found in extant proviruses.

### Structural Conservation of Cas6 Proteins from the Plasmid and Proviral Systems

To investigate whether the Cas6 proteins encoded by the plasmid and proviral CRISPR-Cas systems of *Haloferax lucentense* retain similar structures despite their sequence divergence, we generated AlphaFold2-based S3 models and compared their predicted 3D conformations (Jumper et al. 2021). Structural alignment using ChimeraX (Pettersen et al. 2021) revealed a strong overlap between the two models, particularly across the core RNA recognition and processing domains (Figure 4).

**Figure 4.**
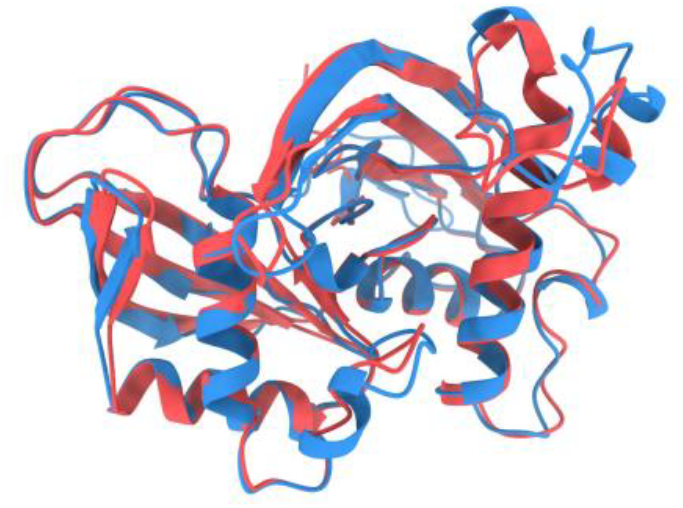
Structural alignment of plasmid-vs provirus-encoded Cas6 from H. lucentense. Overlay of AlphaFold2-predicted 3D models for the Cas6 protein from the megaplasmid-encoded system (shown in blue) and the Cas6 from the proviral system (red). Despite only ∼54% sequence identity, the two structures align almost perfectly (RMSD of 0.777 Å) in their core endoribonuclease fold, including the conserved catalytic pocket. While specific domains are not explicitly labeled in the visualization, the high degree of structural overlap encompasses the regions typically involved in RNA recognition and processing. This near-identical tertiary structure suggests that both Cas6 enzymes perform the same function (pre-crRNA processing) and that strong structural constraints have preserved their fold despite divergent evolutionary origins.

Superposition of the models yielded a backbone RMSD of 0.777 Å over 213 well-aligned residues, with a broader RMSD of 1.957 Å when including all 248 aligned residue pairs. These values indicate a high level of structural conservation, especially within functionally important regions of the protein.

Despite differences in nucleotide composition, codon usage, and even amino acid sequence (as evidenced by ∼54% identity) and differing GC content (63.4% for the proviral system vs 66.8% for the plasmid system), which implies distinct nucleotide composition and codon usage, both viral and plasmid Cas6 proteins adopt a nearly identical tertiary fold. This suggests that their role in pre-crRNA processing is maintained, and that selective pressure may act to preserve protein structure and function even in divergent mobile genetic backgrounds.

Notably, the repeat sequences flanking the CRISPR arrays in the plasmid, proviral, and solitary systems are nearly identical, including exact conservation with repeats found in the *H. volcanii* pHV4 plasmid system. This repeat identity across loci further reinforces the functional compatibility of Cas6 across systems and suggests that, in principle, a single Cas6 enzyme could process crRNAs from all arrays, potentially rendering one of the two encoded Cas6 proteins redundant. This observation adds mechanistic support to the model of functional integration and cross-utilization of CRISPR components within the *H. lucentense* genome (Figure 5).

**Figure 5.**
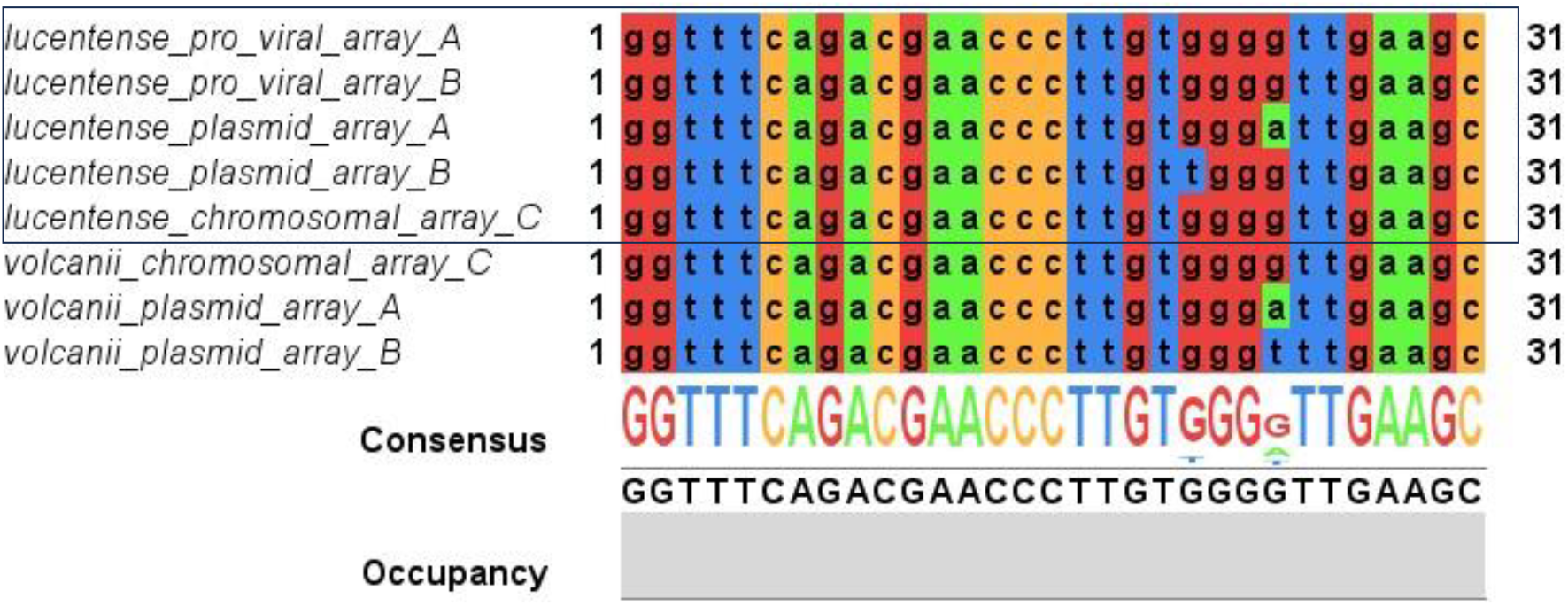
Conservation of CRISPR repeat sequences between Haloferax lucentense and H. volcanii. A multiple alignment of consensus CRISPR repeat units from different arrays highlights nearly identical sequences across species and loci. Repeats from the H. lucentense proviral array and plasmid array match the repeat of the H. volcanii pHV4 plasmid-encoded CRISPR system almost perfectly. Similarly, the solitary chromosomal CRISPR repeat of H. lucentense is identical to the chromosomal repeat in H. volcanii. This high repeat sequence conservation across plasmid, proviral, and chromosomal contexts (and between two Haloferax species) implies a shared evolutionary origin and potential cross-compatibility for spacer acquisition and crRNA processing across systems.

### Differential Selective Pressures on Plasmid and Provirus-Encoded Cas3 Proteins

To investigate the evolutionary pressures acting on the Cas3 proteins, which are crucial for CRISPR interference, we performed a dN/dS analysis using the Selecton server (Stern et al. 2007). This analysis compares the rate of non-synonymous substitutions (dN) to the rate of synonymous substitutions (dS). A dN/dS ratio < 1 indicates purifying selection (pressure to conserve the amino acid sequence), dN/dS ≈ 1 indicates neutral evolution, and dN/dS > 1 indicates positive selection (adaptive evolution favoring amino acid changes). The lower the dN/dS the stronger the purifying selection operating on the gene in question.

This analysis revealed that both the plasmid- and provirus-encoded Cas3 proteins are predominantly under strong purifying selection (dN/dS < >1) across the majority of their sequences. The average dN/dS (ω) ratio for the plasmid Cas3 was 0.190, and for the proviral Cas3 was 0.136, indicating that most amino acid residues in both proteins are under strong functional constraints (Figure 6). This finding aligns with the essential and conserved role of Cas3 in the interference phase of CRISPR-Cas immunity. It is curious in that regard that the selective pressure on the viral systems appears stronger than on the plasmid one in this case. While some sites exhibited elevated ω values, none were supported by confidence intervals indicative of statistically significant positive selection. This suggests that both Cas3 variants are evolutionarily conserved, and there is no strong signal of adaptive diversification in either system. The results imply that, despite their distinct genomic origins and evolutionary trajectories, both Cas3 proteins have retained essential functions required for CRISPR interference and are subject to similar selective constraints.

**Figure 6.**
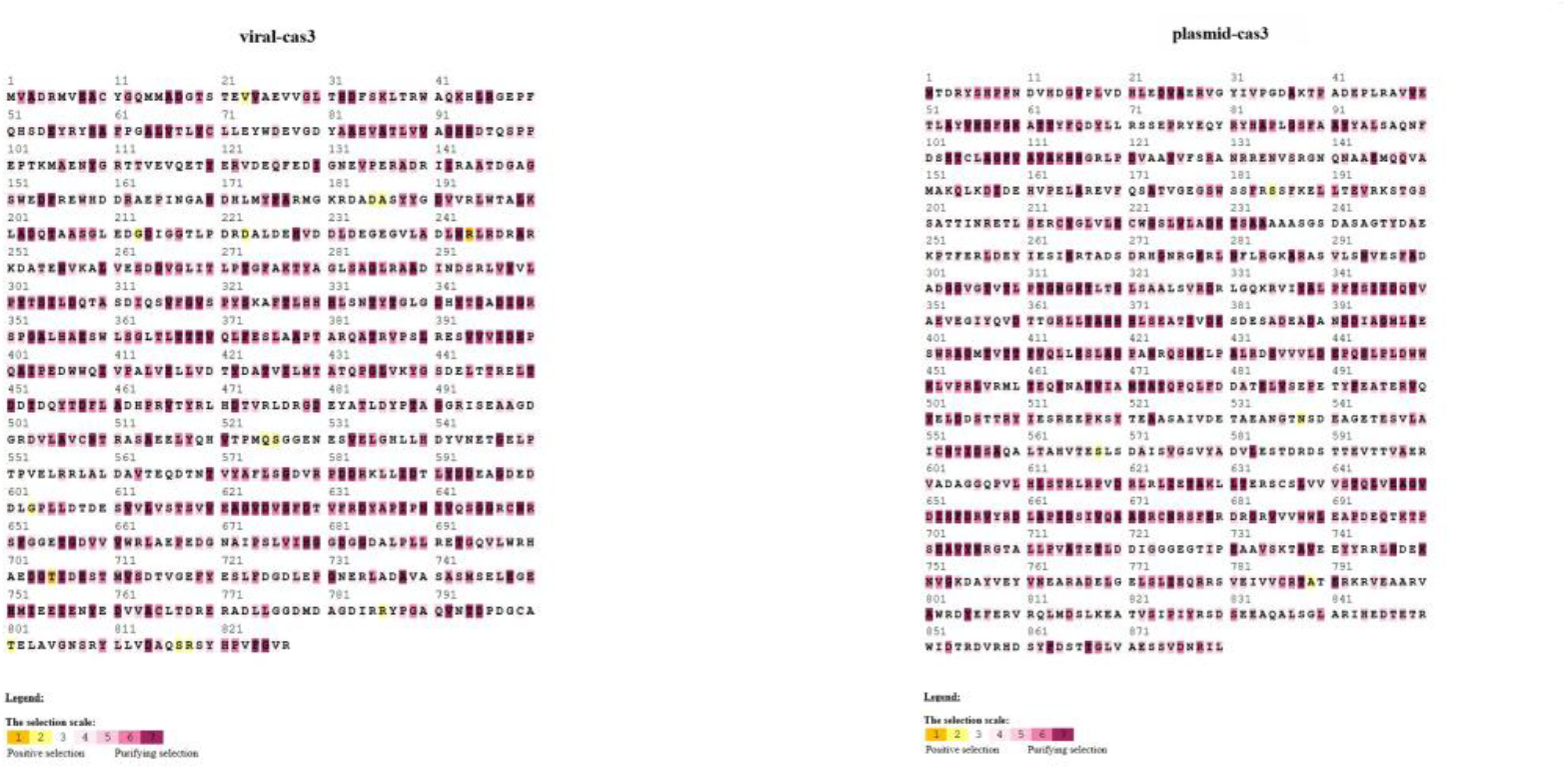
Purifying selection on proviral and plasmid Cas3 proteins at the codon level. Site-by-site dN/dS analysis of the cas3 gene encoding a nuclease/helicase from H. lucentense reveals pervasive purifying selection. Each amino acid position in the Cas3 protein is color-coded by its ω (dN/dS) value (derived from a Selecton analysis): **purple**/dark colors indicate ω ≈ 0 (strong purifying selection), **yellow** highlights sites with ω > 1 (potential positive selection). Panel A shows the provirus-encoded Cas3, and Panel B shows the plasmid-encoded Cas3. Virtually all codons in both Cas3 proteins have ω < 1, with no site exhibiting a statistically significant ω > 1. This indicates that both Cas3 variants are under strong functional constraint, consistent with their essential role in CRISPR interference, and neither shows signs of diversifying (positive) selection.

To assess whether this pattern extends beyond Cas3, we expanded the analysis to include all annotated Cas proteins from both systems. The average dN/dS values for Cas5–8 were also consistently below 1 in both the viral and plasmid systems, reflecting widespread purifying selection across the CRISPR-Cas operons. Among these, Cas3 from the plasmid system exhibited the highest ω value (0.190), suggesting it may be under slightly more relaxed constraint compared to the other Cas proteins. In contrast, Cas7 showed the strongest conservation, with nearly identical ω values of ∼0.071 in both systems, consistent with its role in forming the core backbone of the interference complex (Figure 7). A protein that has to oligomerize and bind additional proteins and crRNA is much more constrained than a protein with higher “freedom”, such as Cas6.

**Figure 7.**
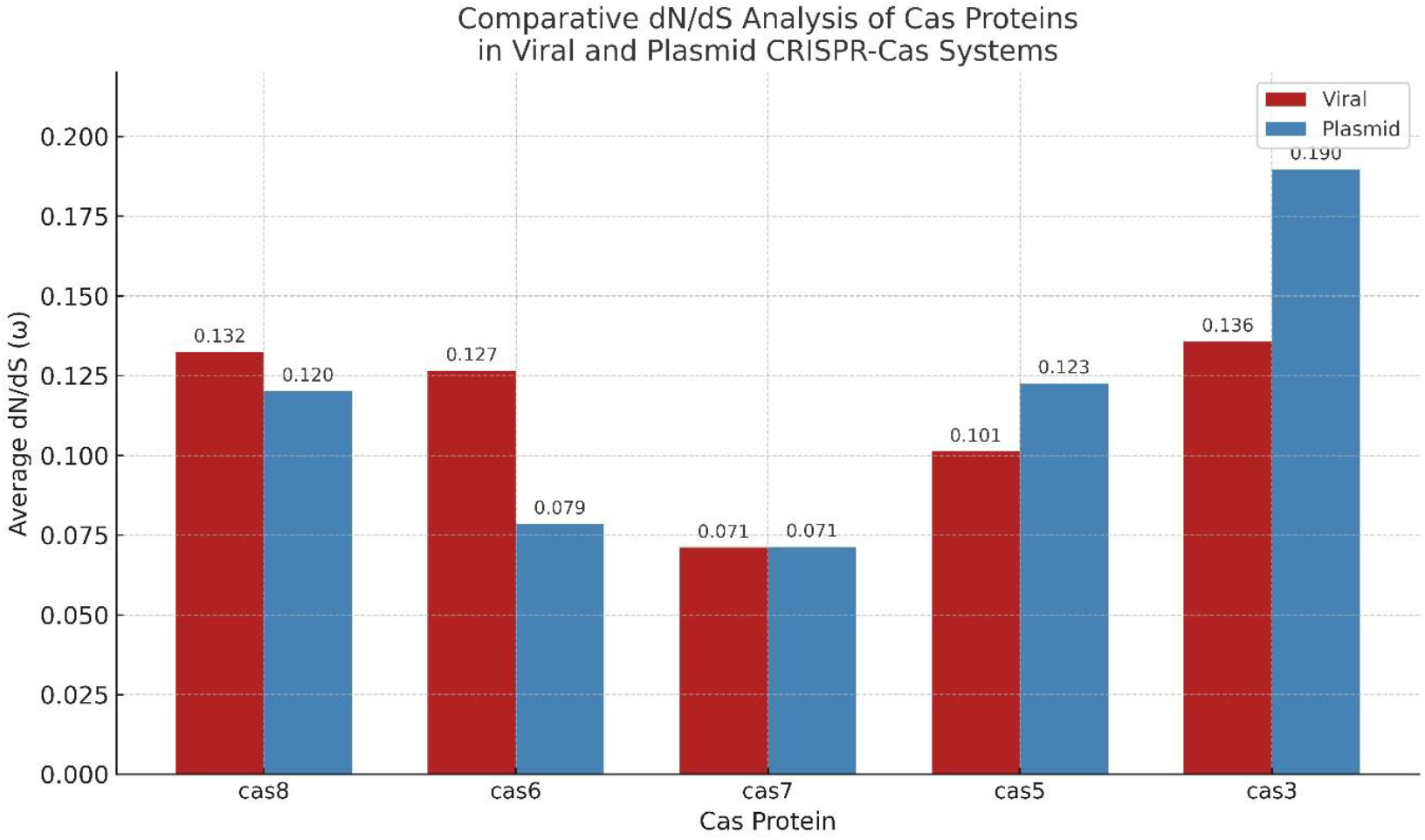
Comparative dN/dS of Cas proteins in proviral vs plasmid CRISPR systems. Comparisons of mean average dN/dS (ω) ratios for each Cas ORF (Cas5 through Cas8, including Cas3 and Cas6) encoded by the proviral systems versus the plasmid systems. All mean ω values are well below 1, indicating purifying selection. Notably, the plasmid-encoded Cas3 has the highest ω (∼0.19) among the set, suggesting it experiences slightly more relaxed selective pressure than the others (though still under purifying selection). In contrast, Cas7 is extremely conserved in both systems, with the lowest ω (∼0.07), fitting its role as a core structural component of the interference complex that must make several functional contacts with proteins and RNA. These trends underscore subtle differences in evolutionary constraint for different Cas proteins, yet overall a strong conservation across the dual systems.

Together, these results support the notion that while all Cas proteins are under purifying selection, there may be subtle differences in evolutionary constraints depending on protein function and genomic context. These data reinforce the idea that despite their mobility and distinct evolutionary origins, both the plasmid- and provirus-encoded systems appear to maintain functionally conserved CRISPR-Cas interference machinery.

## Discussion

Our findings establish *Haloferax lucentense* as a unique archaeal host with an unconventional dual-encoded CRISPR-Cas system, comprising a plasmid-encoded full system and a provirus-encoded incomplete system. The dual system situation also exists in five other genomes, while four genomes only have a viral encoded system, without an additional one (and consequently without the adaptation module and the ability to acquire new spacers). While bioinformatic analyses have revealed distinct evolutionary trajectories, functional modularity, and target specificity, further experimental validation is required to elucidate the precise mechanisms underlying the interaction between these two CRISPR-Cas loci.

The identification of an adaptation-deficient CRISPR-Cas system within a provirus is particularly intriguing. While previous reports have documented partial or defective CRISPR loci in other viruses of archaea and bacteria (Al-Shayeb et al. 2022), our study highlights a functional division of labor between coexisting CRISPR-Cas systems in the same host, raising several key questions. Does the provirus actively leverage the plasmid-encoded adaptation machinery, as suggested by the Black Queen Hypothesis (Morris et al. 2012; Van Valen et al. 1973), or do additional regulatory mechanisms govern the interplay between these systems? The observed spacer overlap between the two systems strongly supports the hypothesis of shared adaptation. The identical CRISPR repeats in the proviral and solitary chromosomal arrays provide a clear mechanism for this: the plasmid’s Cas1-Cas2 complex can recognize these repeats and integrate new spacers into either location (Figure 5).

Another key question pertains to the evolutionary origins and ecological significance of this dual CRISPR-Cas configuration. The presence of nearly identical CRISPR-Cas loci in viruses infecting distantly related haloarchaea suggests horizontal transfer, raising the possibility that viral-encoded CRISPR-Cas systems serve both the conventional function of while primarily contributing to inter-viral competition. This aligns with the concept of “guns for hire,” where defense systems are mobile genetic elements that can be transferred between organisms (Koonin et al. 2020a). The high sequence conservation (>75% nucleotide identity) of the proviral *cas* genes across diverse virus hosts, that often share no similarity in other genes, further suggests that this CRISPR-Cas module is a recent viral innovation that is under strong selective pressure to maintain its functionality, likely for interviral competition.

Why would evolution favor such a split system? The Black Queen Hypothesis (Morris et al. 2012; Van Valen et al. 1973) provides a compelling framework. Adaptation (new spacer acquisition) might be a costly or risky function for a virus to maintain, especially for elements that exist as proviruses and are relatively rarely induced to start replicating. Undesirable unregulated Cas1/Cas2 activity could lead to accidental incorporation of virus self-DNA as spacers, as viruses excise from the chromosome and start replicating (usually using rolling circle replication, a mode previously shown to induce spacer acquisition in archaea (Shiimori et al. 2017)) causing autoimmunity. Having lost *cas1-cas2*, the provirus avoids the danger of misadaptation and relies on the host’s (or in this case, the plasmid’s) better regulated adaptation system, or having no adaptation altogether in those strains that do not encode additional Cas proteins. This “adaptive outsourcing” is not unique; for example, many Type VI CRISPR-Cas systems lack *cas1* and *cas2* and are thought to acquire spacers *in trans* (Silas et al. 2017), and Type IV systems frequently rely on other adaptation modules (Pinilla-Redondo et al. 2022).

The observed target specialization – proviral CRISPR targeting mostly viruses and plasmid CRISPR targeting plasmids – further supports a model of cooperative immunity and division of labor. This pattern is not unique to *H. lucentense*; broader genomic surveys show that plasmid-borne CRISPR loci generally target other plasmids, while chromosomal CRISPR arrays tend to target viruses (Pinilla-Redondo et al. 2022). This suggests that each system in *H. lucentense* is tuned to protect its genetic “owner”: the provirus benefits by eliminating competing viruses, and the plasmid benefits by eliminating competing plasmids. This, in turn, benefits the host cell, by protecting it from being over-populated by costly MGEs.

The structural similarity between the proviral and plasmid-encoded Cas6 proteins, despite their distinct evolutionary origins, demonstrates functional convergence. Both proteins still retain key catalytic residues and predicted active site architectures, ensuring that the proviral system can process crRNAs effectively. This again shows that the provirus has retained the essential components for interference while outsourcing the potentially risky adaptation step.

While our bioinformatic analyses provide strong evidence for the existence of cooperative, dual CRISPR-Cas systems in haloarchaea, it is important to acknowledge the limitations and propose future experimental validation. Although we have shown spacer sharing and target specialization, the precise mechanisms of interaction, such as whether the plasmid’s Cas1-Cas2 complex can access the proviral array and acquire new spacers for it, remain to be demonstrated. Furthermore, while the proviral CRISPR-Cas locus appears to be intact and potentially functional, its activity level and regulation under different conditions need to be determined.

The absence of this system in closely related proviruses highlights the dynamic nature of viral evolution and the potential for horizontal gene transfer to drive the acquisition of novel functions in viruses and other MGEs.

## Materials and methods

### Genomic Sequencing and Assembly

*Haloferax lucentense* strain DSM-14919 was obtained from the DSMZ culture collection (DSM-14919). An initial assembly was downloaded from the NCBI RefSeq database (accession number: GCF_000336795.1) and used for preliminary reference, but not for the final assembly process.

Genomic DNA was extracted from *H. lucentense* cultures using the Qiagen DNeasy Blood & Tissue Kit, following the manufacturer’s instructions. High molecular weight DNA was prepared for Oxford Nanopore sequencing using the Nanopore ligation sequencing kit (Jain et al. 2016). Sequencing was performed across two consecutive Nanopore runs, and the resulting datasets were combined to generate a comprehensive set of long reads. In addition, raw Illumina short reads for *H. lucentense* were downloaded from the NCBI Sequence Read Archive (SRA)(Sayers et al. 2023). Hybrid assembly of the Nanopore long reads and Illumina short reads was performed using the Trycycler pipeline (v0.5.3) with default parameters (Wick et al. 2021b). This approach allowed the generation of a high-quality consensus genome by leveraging the long-read continuity and short-read base accuracy, without relying on the previously available fragmented contigs.

### CRISPR-Cas System Identification and Annotation

Identification of CRISPR-Cas systems in *Haloferax lucentense* and related elements was performed using PADLOC (https://padloc.otago.ac.nz/; Payne et al. 2021), an automated pipeline for detecting and classifying CRISPR-Cas systems. PADLOC output was used to determine the genomic coordinates, predicted system types, and Cas protein compositions for each locus. To assess the genomic context of each CRISPR-Cas system, the surrounding genes were annotated based on the default NCBI GenBank feature tables (Sayers et al. 2022). Annotations of neighboring genes were analyzed to determine whether each CRISPR-Cas system was located on a chromosomal region, a plasmid, or a proviral (phage-derived) element. This contextual annotation was applied both to *H. lucentense* and to related CRISPR-Cas loci identified in other *Haloferax* strains and viral genomes with high sequence similarity to the *H. lucentense* systems.

### Comparative Genomics and Phylogenetic Analysis

Homologs of the Cas proteins identified in *Haloferax lucentense* were retrieved using BLASTP searches (NCBI BLAST+; Camacho et al. 2009). The search database was restricted to the taxonomic group Halobacteria (taxid: 183963; Grant et al. 2001), with the maximum number of target sequences set to 500. This increased retrieval limit was chosen to ensure recovery of both the plasmid-derived and provirus-derived Cas protein homologs, which are highly divergent and would not both be captured using default target limits. BLASTP hits were filtered to retain only sequences with greater than 50% amino acid identity and greater than 95% query coverage relative to the query Cas protein. Redundant sequences were subsequently removed using SeqKit (v0.16.1; Shen et al. 2016), and the resulting non-redundant datasets were aligned using MAFFT (v7.525) with the --localpair and -- maxiterate 1000 options, corresponding to the L-INS-i algorithm optimized for highly accurate local pairwise alignment (Katoh & Standley 2013). Multiple sequence alignments were manually inspected and trimmed using MEGA12 (Kumar et al. 2024) to remove poorly aligned regions and sites containing gaps, thereby improving phylogenetic resolution.

Phylogenetic trees were then reconstructed from these trimmed alignments using the Maximum Likelihood (ML) method implemented in MEGA12 (Kumar et al. 2024). For specific analyses, such as the Cas6 phylogenetic tree which ultimately included 131 amino acid sequences and 186 positions in the final dataset, the Jones-Taylor-Thornton (JTT) amino acid substitution model was employed (Jones et al. 1992). Evolutionary rate differences among sites were modeled using a discrete Gamma distribution with four categories (+G); for the Cas6 tree, the gamma parameter was 0.8751. The robustness of the tree topology was assessed by bootstrap analysis with 100 replicates (Felsenstein 1985). The initial tree(s) for the heuristic ML search were selected by choosing the tree with superior log-likelihood between a Neighbor-Joining (NJ) tree (Saitou and Nei 1987) generated using a matrix of pairwise distances computed using the p-distance (Nei and Kumar 2000), and a Maximum Parsimony (MP) tree (the shortest among 10 MP tree searches, each performed with a randomly generated starting tree). For the Cas6 tree, the analysis yielded a tree with a log likelihood of - 11,154.63. For all ML tree constructions, the option to use all sites present in the final trimmed alignment was selected, and analyses utilized up to 12 parallel computing threads.

The synteny comparison of viral genomes encoding partial CRISPR-Cas systems related to *H. Lucentense* proviral system created (using genebank files of the viruses) with clinker(https://cagecat.bioinformatics.nl/tools/clinker; Gilchrist et al. 2021)

### Spacer Analysis and Target Prediction

CRISPR arrays and associated spacer sequences were identified using the CRISPRCasFinder tool available at https://crispr.otago.ac.nz/ (Couvin et al. 2018). Spacer target prediction was performed using CRISPRTarget (https://crispr.otago.ac.nz/CRISPRTarget) against the default MGE and viral database provided on the platform. For target prediction, the E-value threshold was set to 0.01 (more stringent than the default threshold of 1) to improve specificity and minimize false-positive hits. For each spacer with a predicted target, the corresponding target accession was manually examined to determine the genomic context. Neighboring gene annotations were analyzed to classify the target as being located on a plasmid, viral genome, provirus, or chromosomal region, and to infer potential target function based on adjacent gene features. The table of targets and all spacers available in the supplementary_data_3 file.

### Protein Structure and Function Analysis

Predicted protein structures for the Cas6 proteins encoded by the plasmid and proviral CRISPR-Cas systems of *Haloferax lucentense* were retrieved from the AlphaFold Protein Structure Database (https://alphafold.ebi.ac.uk/) (Jumper et al. 2021). The amino acid sequences of the Cas6 proteins were used as queries, corresponding to UniProt accessions A0A6C0UUT1 (proviral Cas6) and M0GYZ3 (plasmid Cas6). The resulting predicted structures (PDB format) were downloaded for further comparative analysis. Structural comparison and alignment of the plasmid- and provirus-derived Cas6 models were performed using UCSF ChimeraX (latest version available at the time of analysis). The two models were superimposed using the matchmaker command (mmaker #1 to #2), aligning based on backbone alpha-carbon atoms to assess structural similarity.

### CRISPR Repeat Sequence Alignment

Consensus repeat sequences were extracted from the CRISPR arrays of *H. lucentense* (proviral, plasmid, and solitary chromosomal arrays) and from the corresponding CRISPR loci in *Haloferax volcanii*, including the pHV4 plasmid and chromosomal arrays. These sequences were aligned using Jalview version 2.11.4.1 with default alignment settings (Waterhouse et al. 2009). The alignment was visualized using Jalview’s color-coded residue scheme to highlight conserved nucleotide positions and variations. The consensus sequence and occupancy bar were used to assess repeat conservation across arrays.

### Selection Regime Analysis

To evaluate the evolutionary constraints acting on the Cas proteins encoded by the proviral and plasmid CRISPR-Cas systems of *Haloferax lucentense*, we conducted a codon-level analysis of selection pressure using the Selecton server (http://selecton.tau.ac.il/) (Stern et al. 2007). The goal was to compare the strength and distribution of purifying versus positive selection across all core Cas components in both systems. For each Cas protein (Cas3, Cas5, Cas6, Cas7, Cas8), BLASTP searches were performed against the NCBI nr database using the *H. lucentense* protein sequences as queries.

Homologs showing ≥75% amino acid identity were selected, and their corresponding coding DNA sequences (CDS) were manually retrieved from NCBI. Redundant entries (e.g., identical sequences from the same or closely related strains) were removed manually to retain one representative per unique allele. Each curated CDS dataset were submitted to the Selecton web server using its default parameters, which allows for variable selection across sites (ω < 1 for purifying selection; ω > 1 for positive selection). For visualization and interpretation, color-coded selection maps were generated by Selecton for each Cas gene (supplementary_data_2).

## Supporting information

Supplementary dataset 2 - Selecton data

Supplementary dataset 1 - phylogenetic trees

Supplementary dataset 3 - spacer data

## Data Availability

The assembled genome of *H. lucentense* is included in the supplementary_data_3 file. Accession numbers for all genomes and the specific proviral regions analyzed are also provided in the supplementary materials. Scripts used for custom analyses are available upon request.

## Acknowledgements

The authors thank Neta Altman for a critical reading of the manuscript. This work was funded by a European Research Council (grant ERC-AdG 787514) and the DFG priority programme “CRISPR-Cas functions beyond defence” SPP2141 grants to UG.

