## Supplementary figures and images for "Evolutionary insights into provirus-encoded CRISPR-Cas systems in halophilic archaea"

### cas3_final_tree.pdf

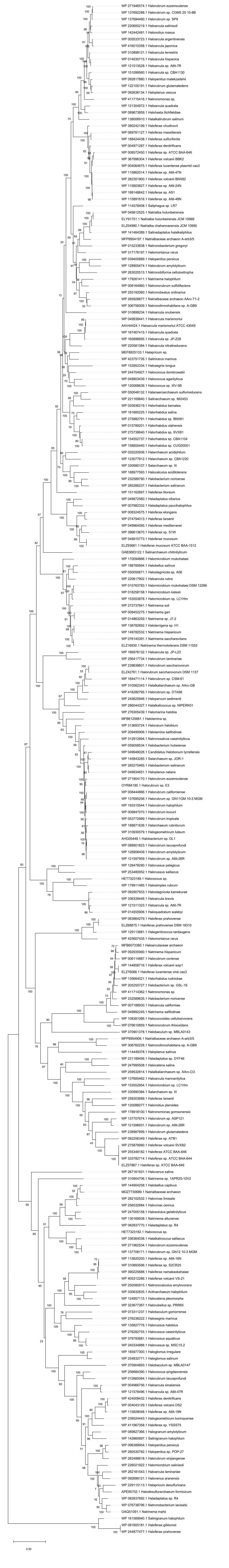

### cas5_final_tree.pdf

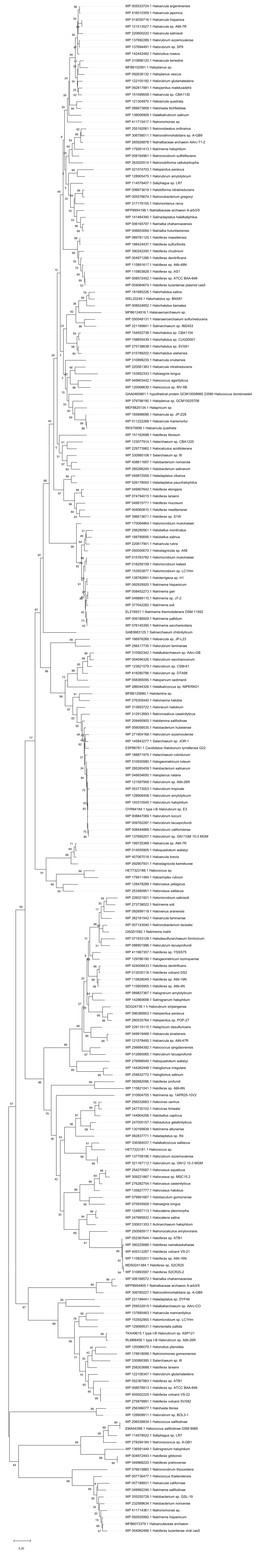

### cas5_plasmid_resutls_selecton_final.png

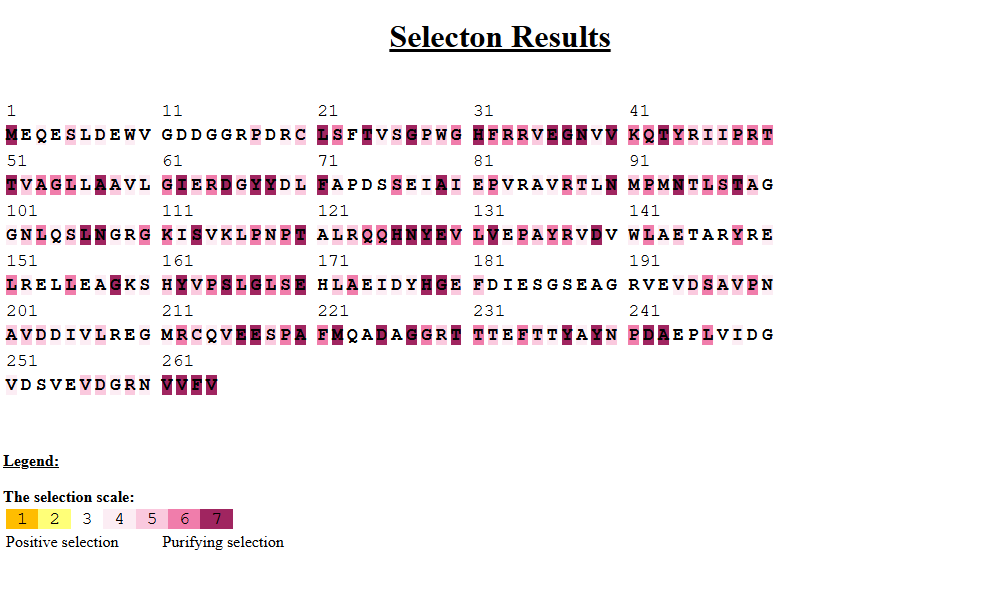

### cas5_viral_final_selecton.png

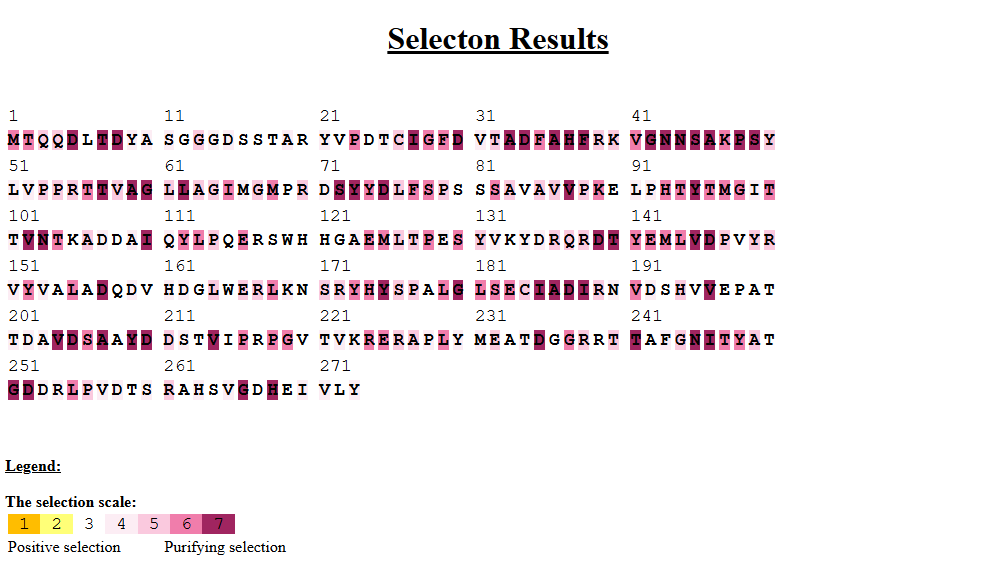

### cas6_final_tree.pdf

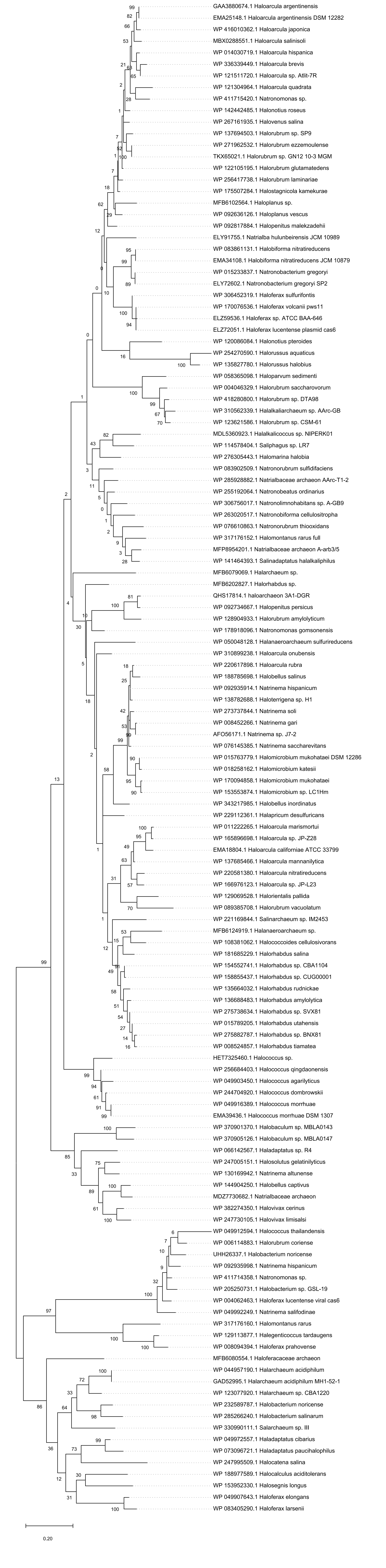

### cas6_plasmid_results_selecton_final.png

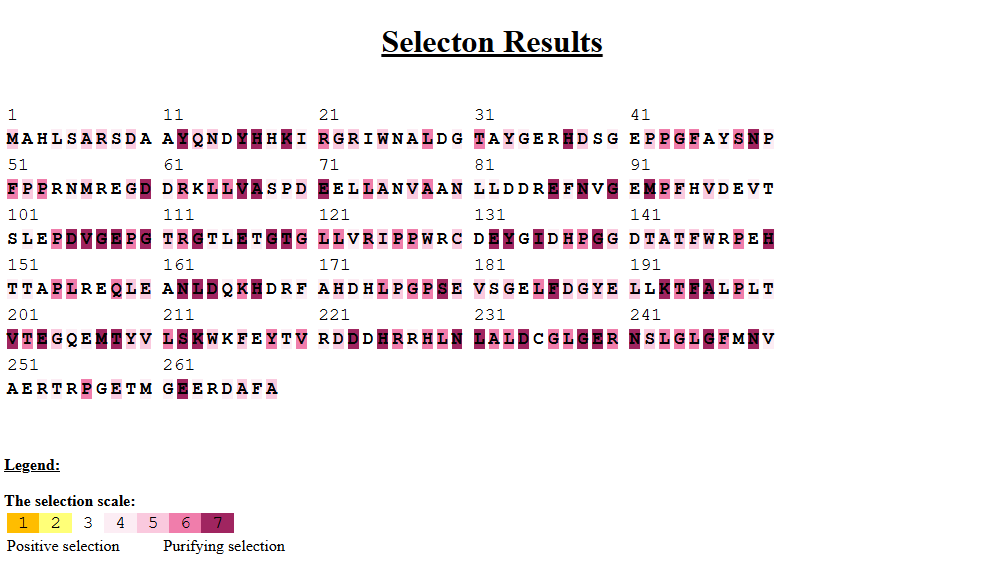

### cas6_viral_selecton_final.png

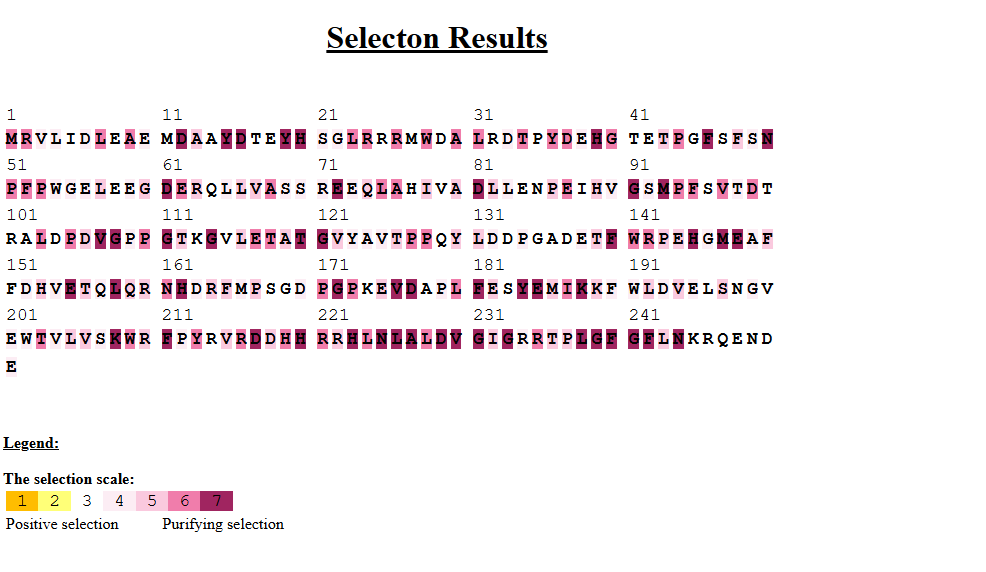

### cas7_final_tree.pdf

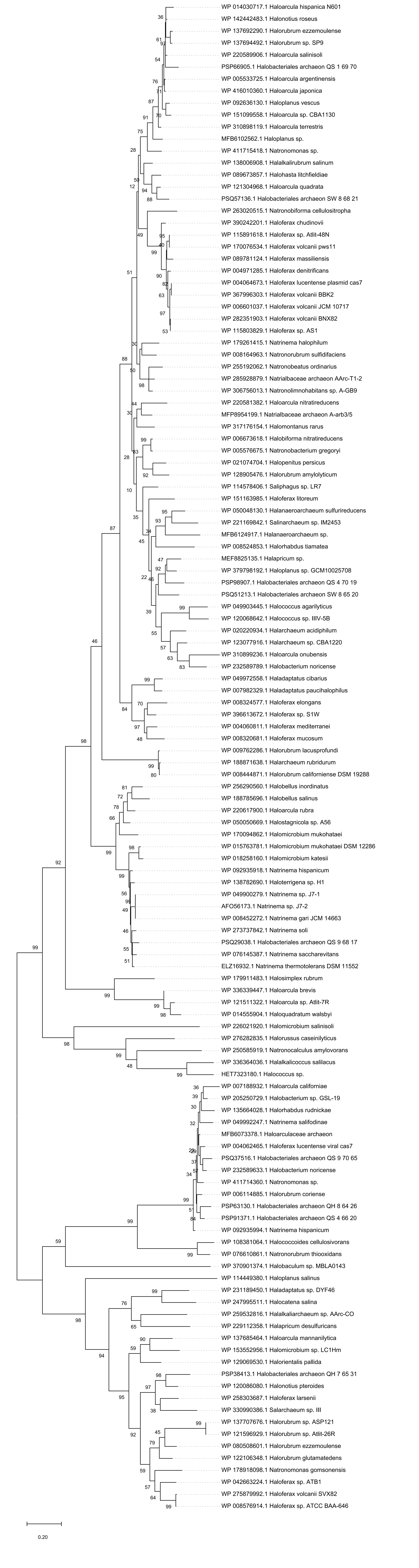

### cas7_plasmid_results_selecton_final.png

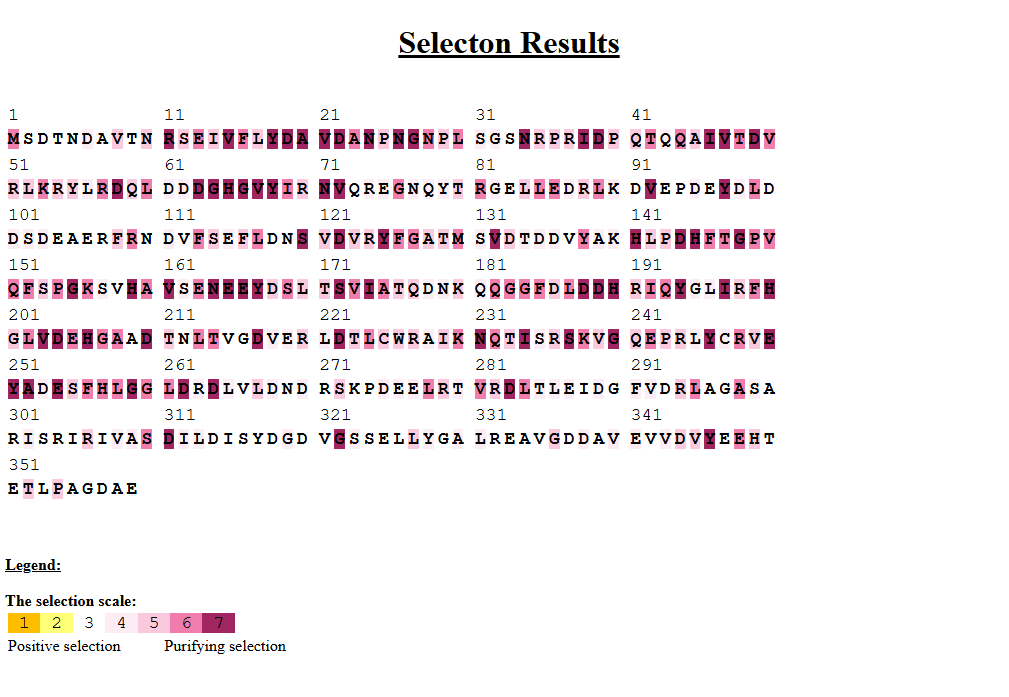

### cas7_viral_final_selecton.png

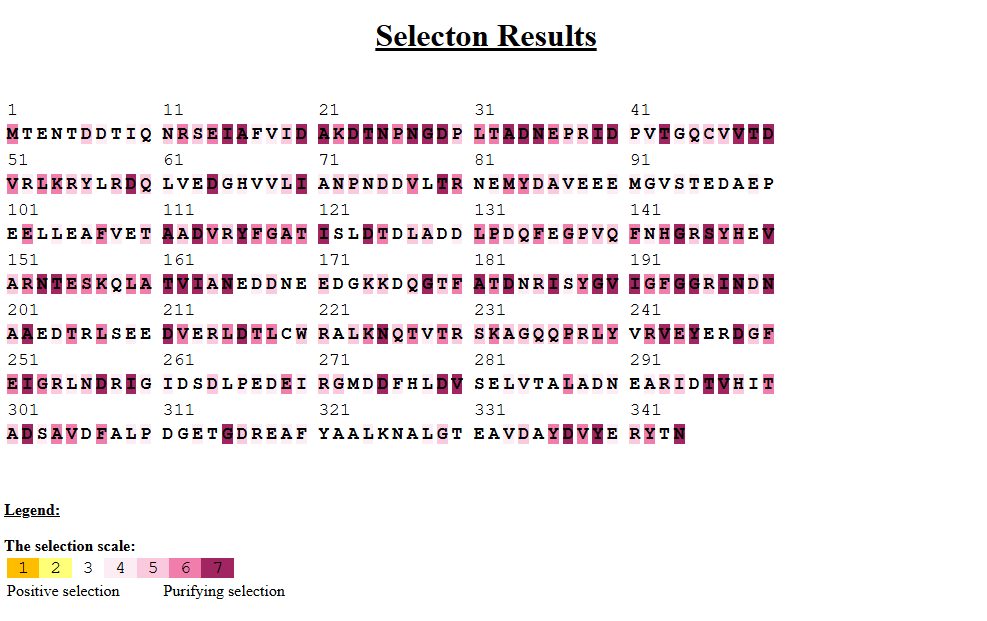

### cas8_final_tree.pdf

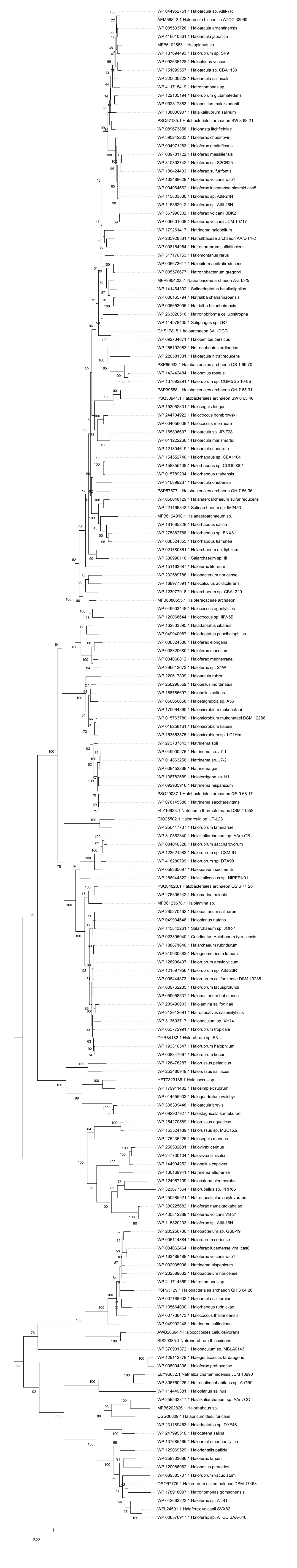

### cas8_plasmid_resutls_selecton_final.png

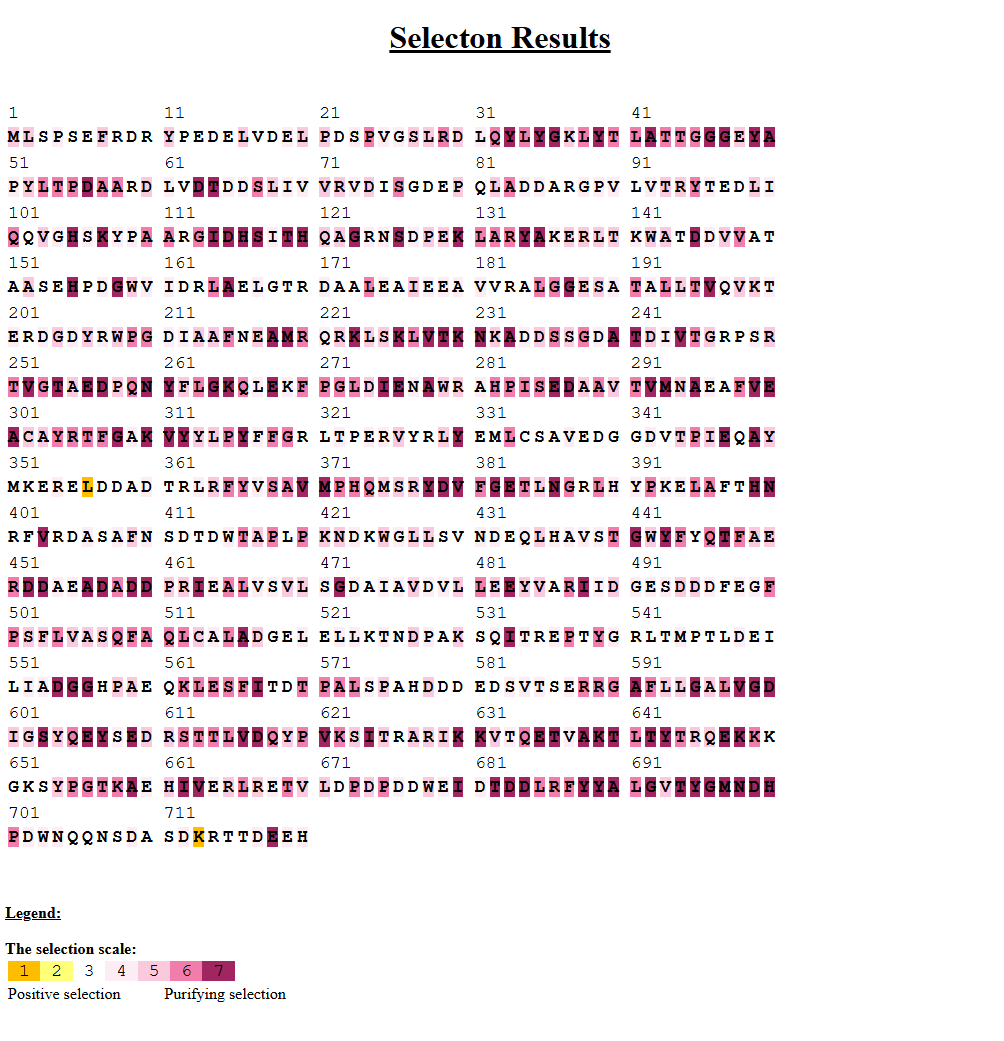

### cas8_viral_selecton_final.png

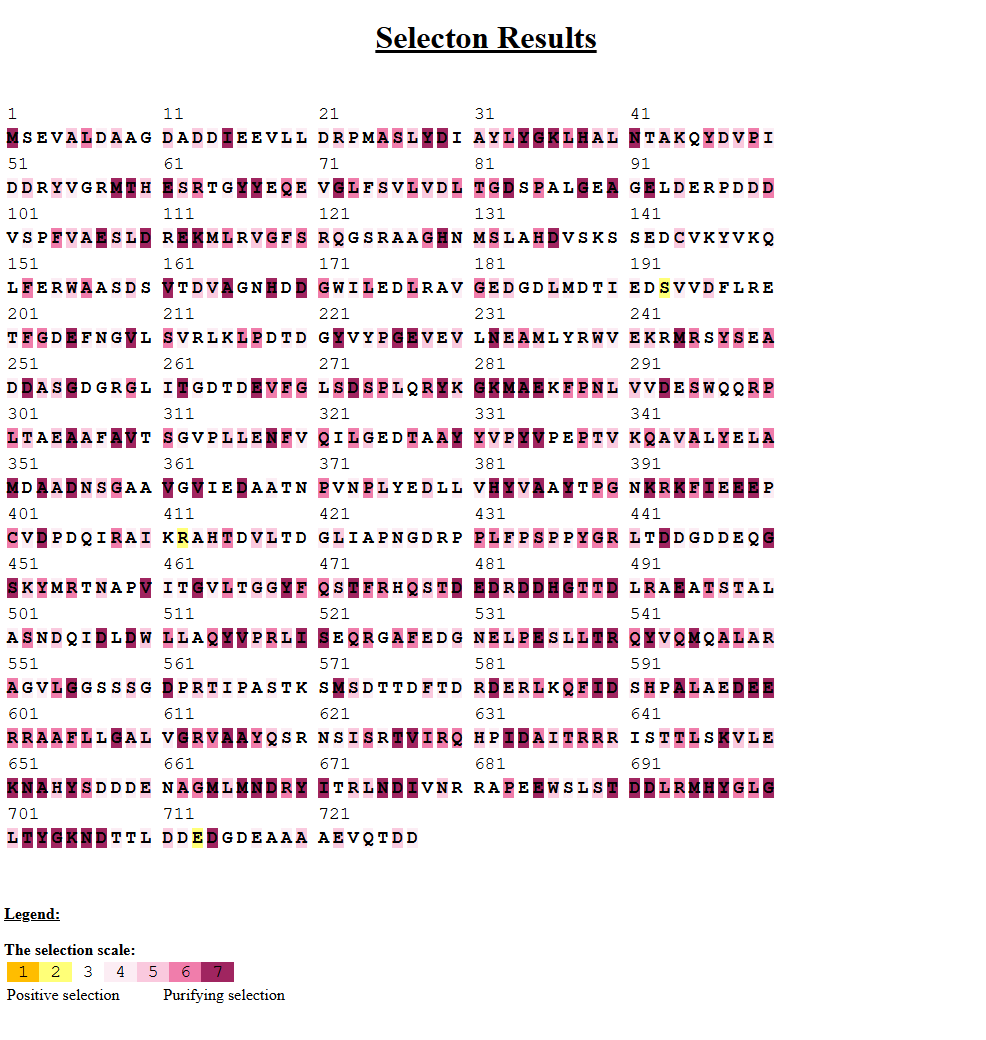

### plasmid_cas3.png

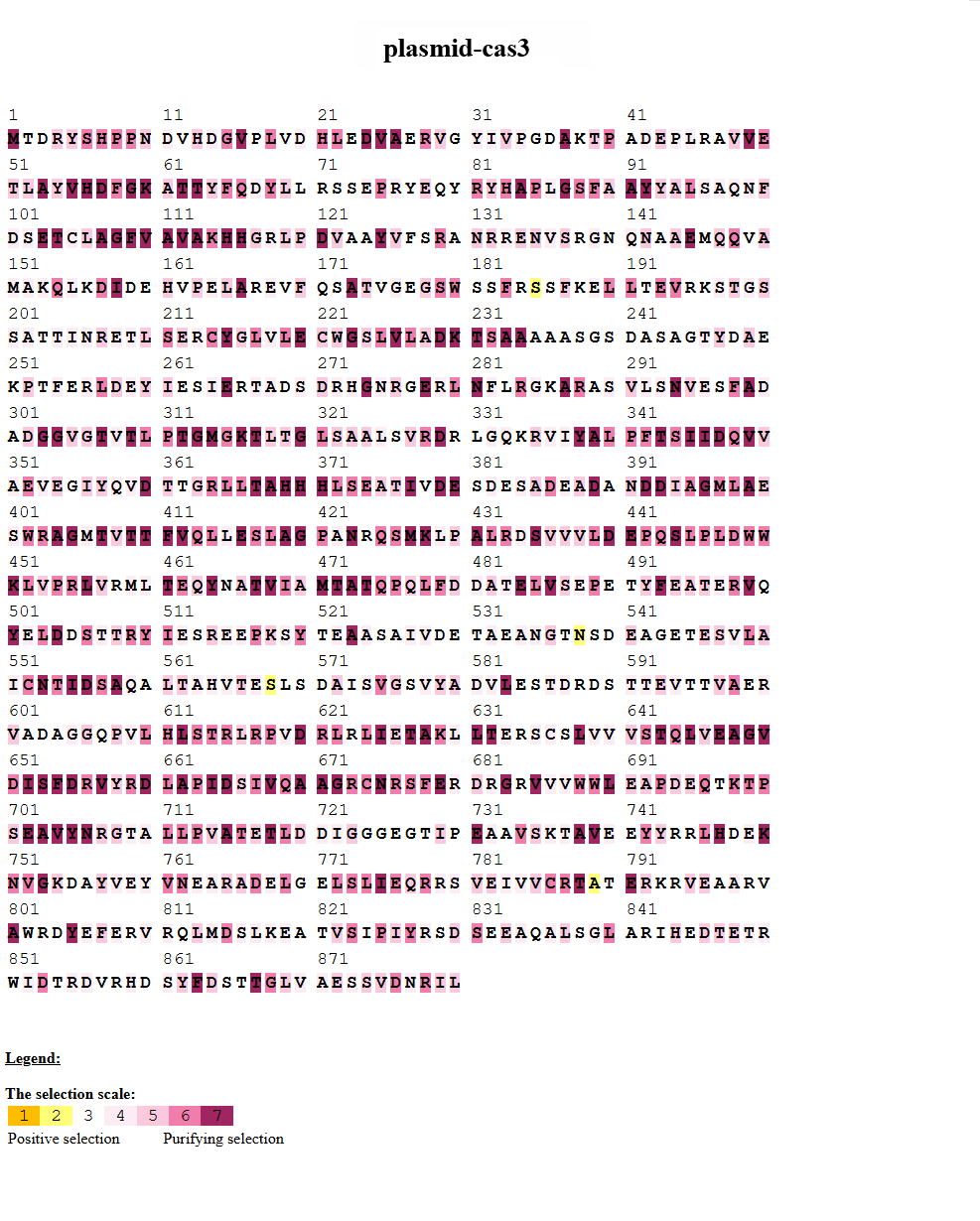

### viral_cas3.png

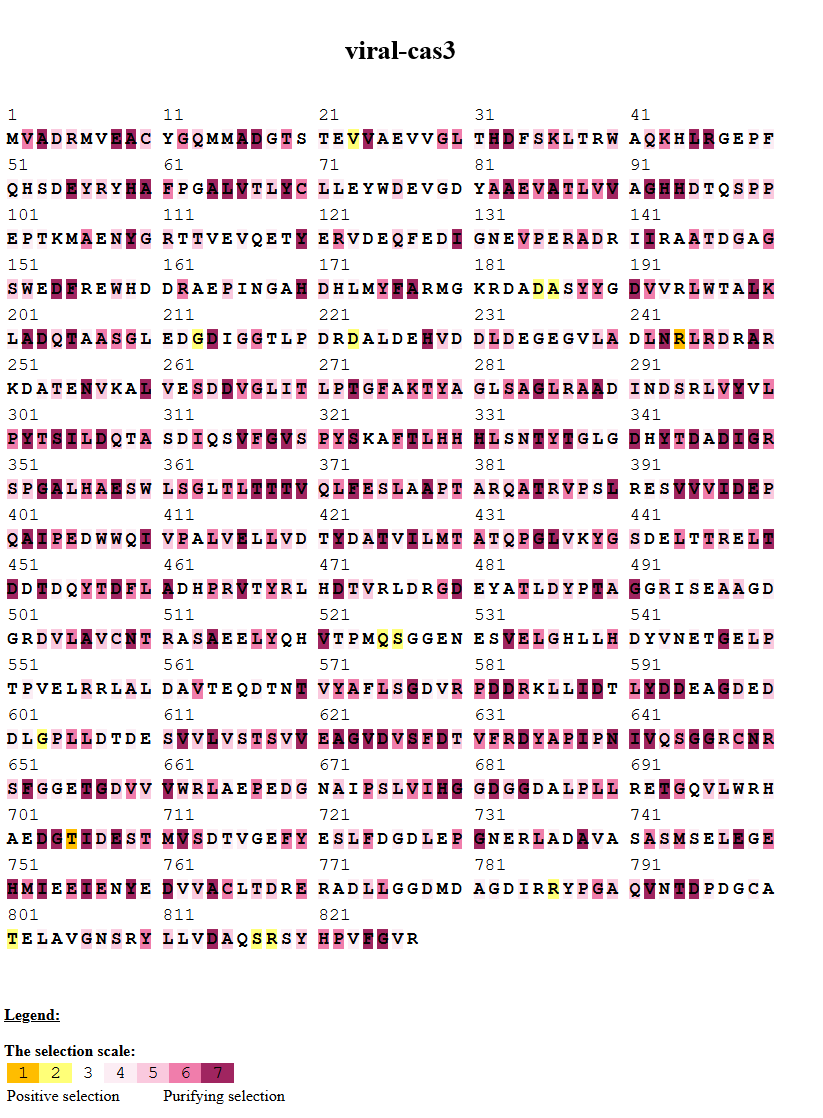
